# Application of t-SNE to Human Genetic Data

**DOI:** 10.1101/114884

**Authors:** Wentian Li, Jane E Cerise, Yaning Yang, Henry Han

## Abstract

The t-SNE (t-distributed stochastic neighbor embedding) is a new dimension reduction and visualization technique for high-dimensional data. t-SNE is rarely applied to human genetic data, even though it is commonly used in other data-intensive biological fields, such as single-cell genomics. We explore the applicability of t-SNE to human genetic data and make these observations: (i) similar to previously used dimension reduction techniques such as principal component analysis (PCA), t-SNE is able to separate samples from different continents; (ii) unlike PCA, t-SNE is more robust with respect to the presence of outliers; (iii) t-SNE is able to display both continental and sub-continental patterns in a single plot. We conclude that the ability for t-SNE to reveal population stratification at different scales could be useful for human genetic association studies.

## Background

Genome-wide association study (GWAS) [1, 2, 3, 4, 5, 6] is an approach to type common genetic variants in a general human population, and by comparing the allele frequency difference between a group of patients (cases) and a group of normal people (controls), discover statistical association signals. The genetic variant mostly encountered is the single nucleotide polymorphism (SNP) [7]. These associated variants may reside in a protein coding region, indicating a possible change of transcription products which may play a role in the disease. It may also sit between genes which probably change a binding motif of a transcription factor which leads to a change of the transcription (expression) level [8]. A comprehensive database on statistically significant variants for many human diseases is the GWAS Catalog, first maintained by the NHRGI/NIH (National Human Genome Research Institute of National Institute of Health (USA)), then by the European Bioinformatics Institute (UK), which can be found at *https://www.ebi.ac.uk/gwas/* [9, 10].

A key step in GWAS analysis is to match the ethnicity background of cases and controls. Failure to do so would confound allele frequency difference due to disease-causing mutations and difference due to population genetic history. There have been many attempts to correct this “spurious association” due to “population stratification” [11, 4]: “genomic control” uses non-functional variants to estimate the amount of population genetic history differences and the association signal is corrected accordingly [12]; “family-based association” uses untransmitted alleles as controls thus circumventing the population stratification issue completely when these type of data are available [13, 14, 15]; K-mean clustering to group sample genotype data towards the presumed K populations [16]; incorporating co-variance among samples to correct the association signal [17, 18, 19], etc.

However, the most common practice in dealing with population stratification or other subtle/hidden structures in genetic data is to perform dimensional reduction techniques, such as MDS (multi dimensional scaling), PCA (principal component analysis), SVD (singular value decomposition). The reduced dimensions can be directly visualized and, in the case of PCA, can be used as covariates in the association analysis [20, 21, 22, 23, 24, 25, 26]. Within the European populations, it is well established that the first PC (PC1) aligns with the north-south direction (latitude), and the second PC (PC2) aligns with the east-west direction (longitude) [27].

Though the use of PCA is mostly satisfactory, the method is not without a problem. Most notably, PCA is highly affected by the presence of outliers. If most of the samples belong to one homogeneous population with a minority from another different population, the presence of genetically different minority samples can completely change the principal axes, thus changing the distribution of samples along the main PCs. This is because as a linear holistic dimension reduction method, PCA cannot capture local data characteristics well. Other problems include the determination of the number of SNPs to be included, the role common vs. rare variants play in the result, and the number of PCs to be kept.

Here we explore the application of a new dimensional reduction technique, t-SNE (t-distributed stochastic neighbor embedding) [28], to the genetic data. Similar to MDS, the aim of t-SNE is to preserve the pairwise distance in high-dimensional space to 2 or 3 lower dimensions. Unlike MDS or PCA, the preservation in t-SNE is non-linear: t-SNE minimizes the Kullback-Leibler divergence between two distributions – one distribution that measures pairwise similarities of input samples in high-dimensional space, another heavy-tailed Student's *t*-distribution that measures pairwise similarities of corresponding samples in the lowdimensional embedding space. t-SNE has demonstrated its built-in advantages in capturing local data characteristics and revealing subtle data structures in visualization, as shown in the original publication [28].

t-SNE is a popular choice in the analysis of single-cell RNA-seq data (e.g. [29, 30, 31]), but has not been applied extensively to genetic data. In the single previous publication where t-SNE was applied to genetic data [32], the main conclusion in is that if t-SNE is considered as a clustering technique, it performs better than PCA. We are particularly interested in t-SNE's claimed ability to “reveal structure at many different scales” [28], as major population stratification co-exists with other small-scaled shared evolutionary history among samples.

## Results

### Continental separation

We first examine the ability of t-SNE to separate continental populations (Africa, Asia, Europe, etc.). GWAS usually refers to genetic association studies using common variants. The new next-generation-sequencing (NGS) technique, though aiming at typing all variants, also produces common variants. We use one of the major public NGS data, the 1000 Genomes Project [33, 34, 35]. We extracted 3825 common variants which pass a quality control (QC) criteria, and are also present in the Illumina Global Screening Array chip. We use the KING program to calculate the inter-sample genetic distances. We override the default setting of KING to retain only 3 significant digits so that 9 significant digits are kept, in order to improve our ability to distinguish subtle structures.

Fig. 1 shows the results from PCA (top) and t-SNE (bottom) with the first and the second major dimensions (left column) and the second & third dimension. The number of samples in each continent (though Asia is split into east and south Asia) are more or less balanced: 661 Africans (AFR), 347 “Americans” (AMR), 504 East Asians (EAS), 489 South Asians, and 503 Europeans (EUR). Note that some subgroups living in continental America is not grouped with AMR: African-American in Southwest of US (ASW) and African-Caribbean in Barbados are in the AFR group, Utah CEPH families are in the EUR group, etc.

**Figure 1:**
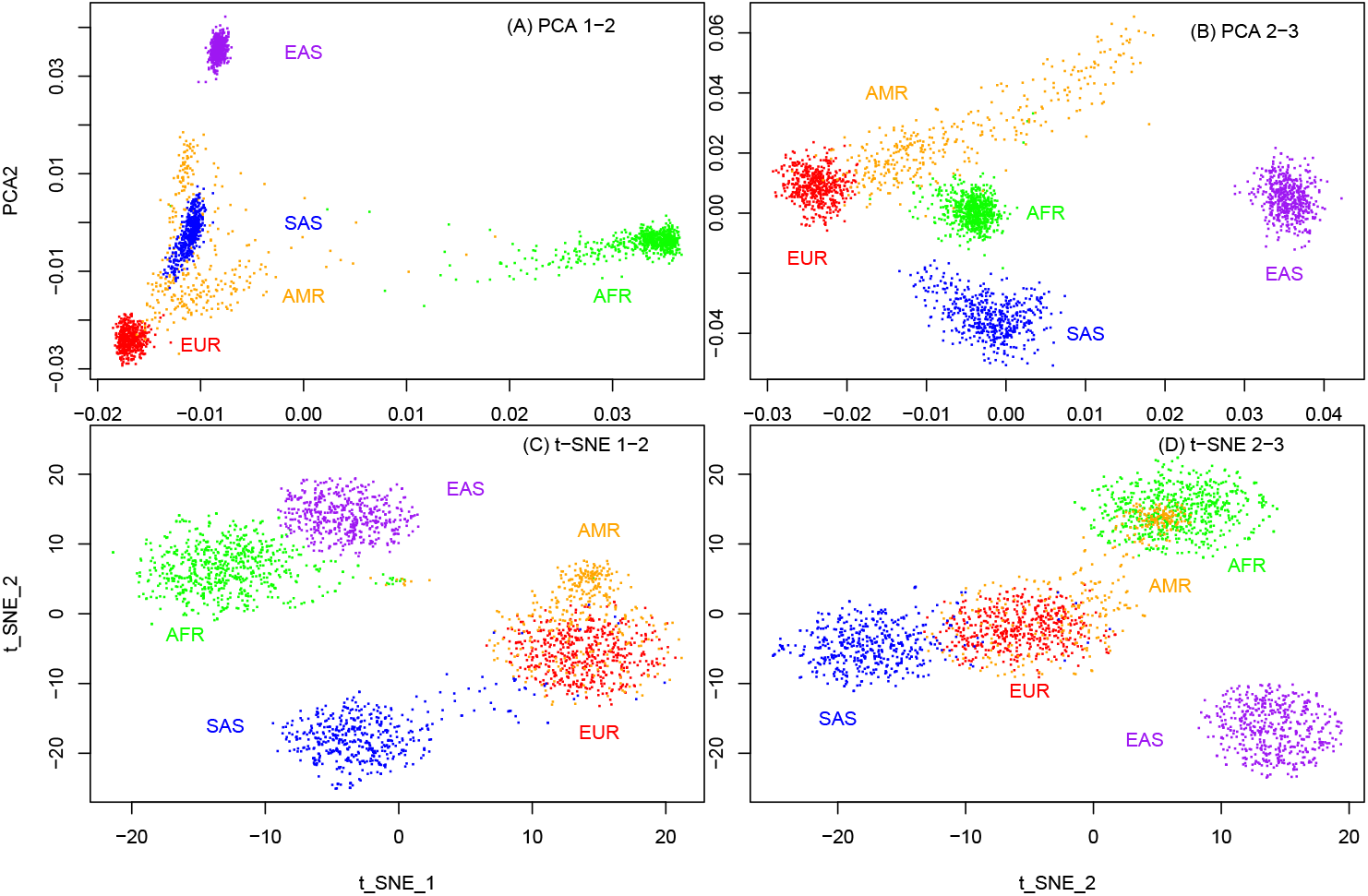
A comparison of three different low-dimensional display of 1000 Genomes Projects data with 3825 SNPs. The number of samples (dots) is 2504, with 661 AFR (African green), 347 AMR (American, orange), 504 EAS (East Asia, purple), 489 SAS (South Asia, blue), and 503 EUR (European, red). (A) PCA (PC1-PC2); (B) PCA (PC2-PC3); (C) t-SNE (1-2); (D) t-SNE (2-3).

Although all methods are able to separate continental populations, PCA (1-2 dimensions) shows an overlap between South Asian and American, whereas t-SNE shows AMR has more overlap with Europeans. In the 2-3 dimension, PCA shows some link between AMR and EUR, whereas t-SNE continue to show a strong connection between AMR and EUR. As some AMR samples, such as those from Colombia, Puerto Rico, and to some extent, Mexico (actually the samples are Mexican-American), are expected to contain European ancestor, the t-SNE result is consistent with our external knowledge. As can be seen in the 3D version of t-SNE (Fig. 2), the overlap between AMR and AFR in Fig. 1(F) is an artifact, as AMR and AFR are actually separated.

**Figure 2:**
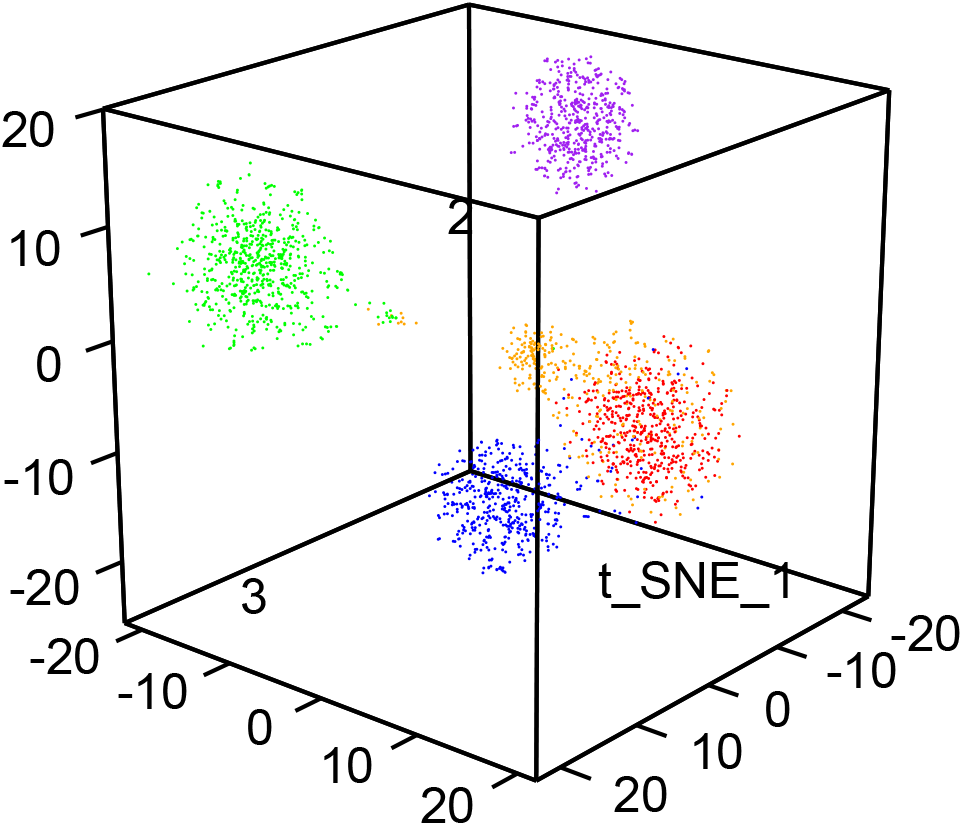
3-dimensional t-SNE which combines information from Fig. 2(E) and (F). Color scheme: green for AFR, orange for AMR, purple for EAS, blue for SAS, and red for EUR.

### Treatment of outliers

We compare t-SNE and PCA in a more realistic setting where most of the samples belong to one ethnic group, whereas a few are either from distinct ethnic groups or a mixed race. For that purpose, we extract 99 Utah residents with North/West European ancestry (CEU), 91 England/Scotland samples (GBR), and 5 African-American in Southwest (ASW), 5 Mexican-American in Los Angeles (MXL), 2 Chinese in Beijing (CHB), 2 Chinese in South China (CHS).

PCA (Fig. 3 (A)) moves ASW, CHB/CHS far away from the main cluster of roughly 200 Caucasians, whereas MXL samples, though still separated, are closer to the center. However, though t-SNE (Fig. 3 (B)) shows ASW, CHB/CHS as separated groups, the MXL samples are much closer to other Caucasian samples. In order to see the structure within the Caucasian group through PCA, the usual procedure is to remove the “outliers” and re-run PCA again (e.g. recommended in smartpca). On the other hand, an advantage of t-SNE is to show both the “outliers” and the main cluster with detail simultaneously, capturing subtle local data structures and preserving global structures in the low-dimensional space.

**Figure 3:**
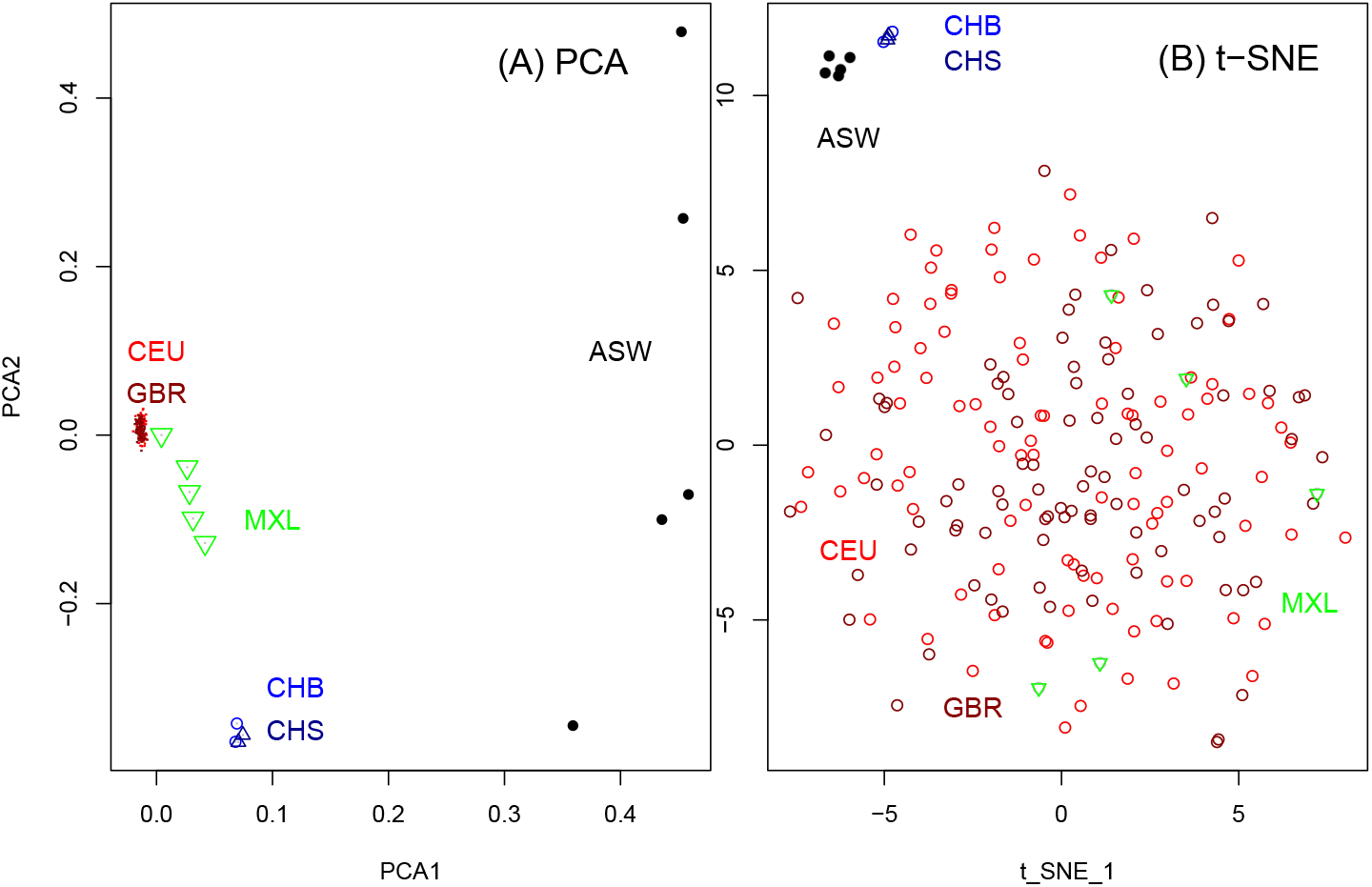
A comparison of PCA and t-SNE on a mostly Caucasian dataset (99 Caucasians from Utah (CEU), USA and 91 Caucasians from UK (GBR)) with “outliers” included (5 African-American in Southwest (ASW), 5 Mexican-American in Los Angeles (MXL), 2 Chinese in Beijing (CHB), 2 Chinese in South China (CHS)). (A) PCA (PC1-PC2); (B) t-SNE.

To see whether some PCA modifications may exhibit similar properties as t-SNE, we test two PCA variants: non-negative PCA (NPCA) [36, 37] and derivative component analysis (DCA) [38]. The purpose of imposing extra constraints in NPCA is that positive and negative terms in classic PCA may cancel each other, leading to a loss of local feature. The purpose of DCA is to use derivatives to capture latent patterns and to suppress noise level. Fig.4 and 5 shows the 3D NPCA and DCA plot. Although NPCA and DCA are better than classic PCA in showing more spreading in CEU/GBR cluster, they are more consistent with classic PCA than with t-SNE.

**Figure 4:**
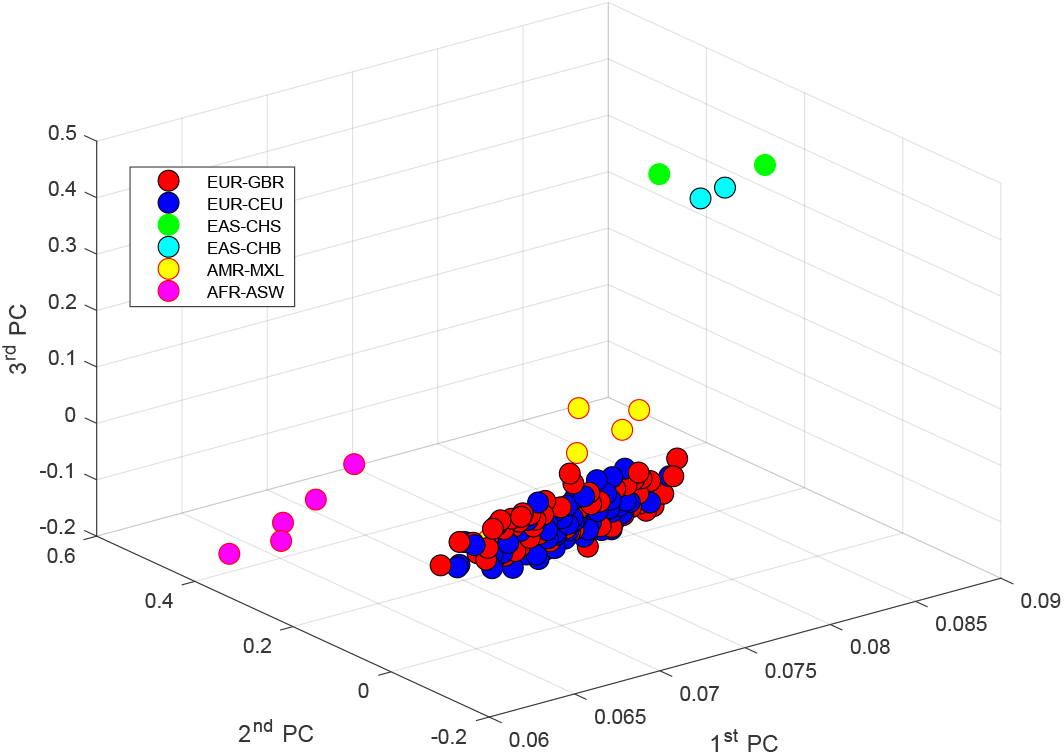
Non-negative PCA (NPCA) of the data used in Fig.3.

**Figure 5:**
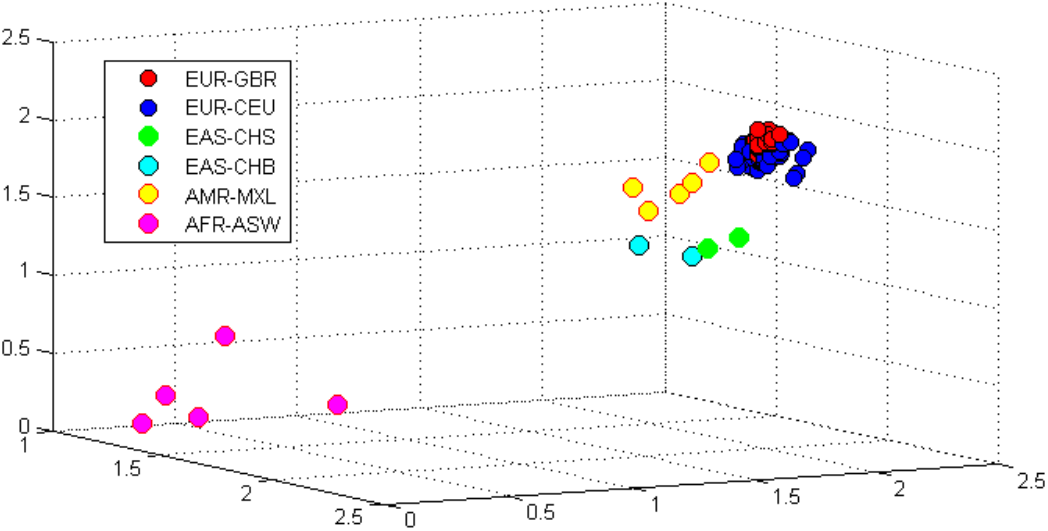
Derivative component analysis (DCA) of the data used in Fig.3.

### Sub-continental separation

We examine whether t-SNE is able to display sub-continental patterns besides the continental separation. For that, we extract a denser set of SNPs from chromosome 1 in the 1000 Genomes Project with the criteria of alternative/minor allele frequency > 0.2, and spacing between neighboring SNPs > 20, 000 bases. This leads to more than 9000 SNPs, roughly equivalent to a genome-wide +100,000 SNPs. The use of a single chromosome to represent the whole genome is justified, as PCAs based on any human chromosome are almost identical (Fig.S3 of [39]). Only in the extreme case of using SNPs from a region with inversion, may the shape of PCA be different [40, 39], as the trimodal distribution of the samples reflects the three underlying configurations [41].

Fig.6 shows the t-SNE with a finer group labeling. We paid particular attention to choose colors to represent a population's geographic information within the continent, which is shown in Fig.7. Generally, we choose a dark color for the population towards the north, and a light color for those in the south. We also switch the t-SNE-1 and t-SNE-2 so that the north groups point to the up direction.

**Figure 6:**
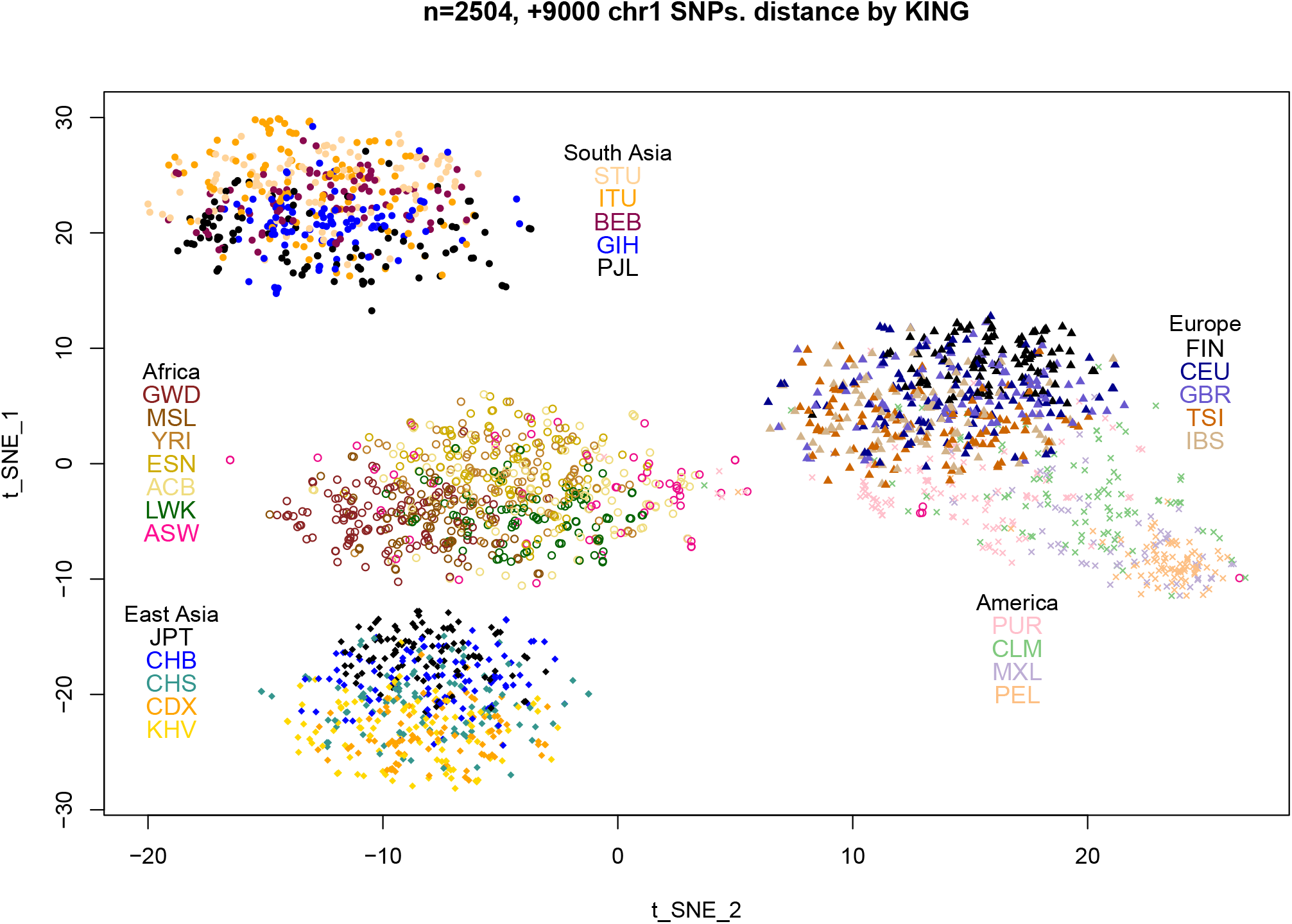
t-SNE plot obtained from more than 9000 SNPs on chromosome 1 of the 1000 Genome Projects (alternative allele frequency > 0.2, spacing between neighboring SNPs longer than 20,000 bases. The explanation of group labels in Africa (ACB, ASW, ESN, GWD, LWK, MSL, YRI), America (CLM, MXL, PEL, PUR), East Asia (CDX, CHB, CHS, JPT, KHV), South Asia (BEB, GIH, ITU, PJL, STU), Europe (CEU, FIN, GBR, IBS, TSI) are explained in the text.

**Figure 7:**
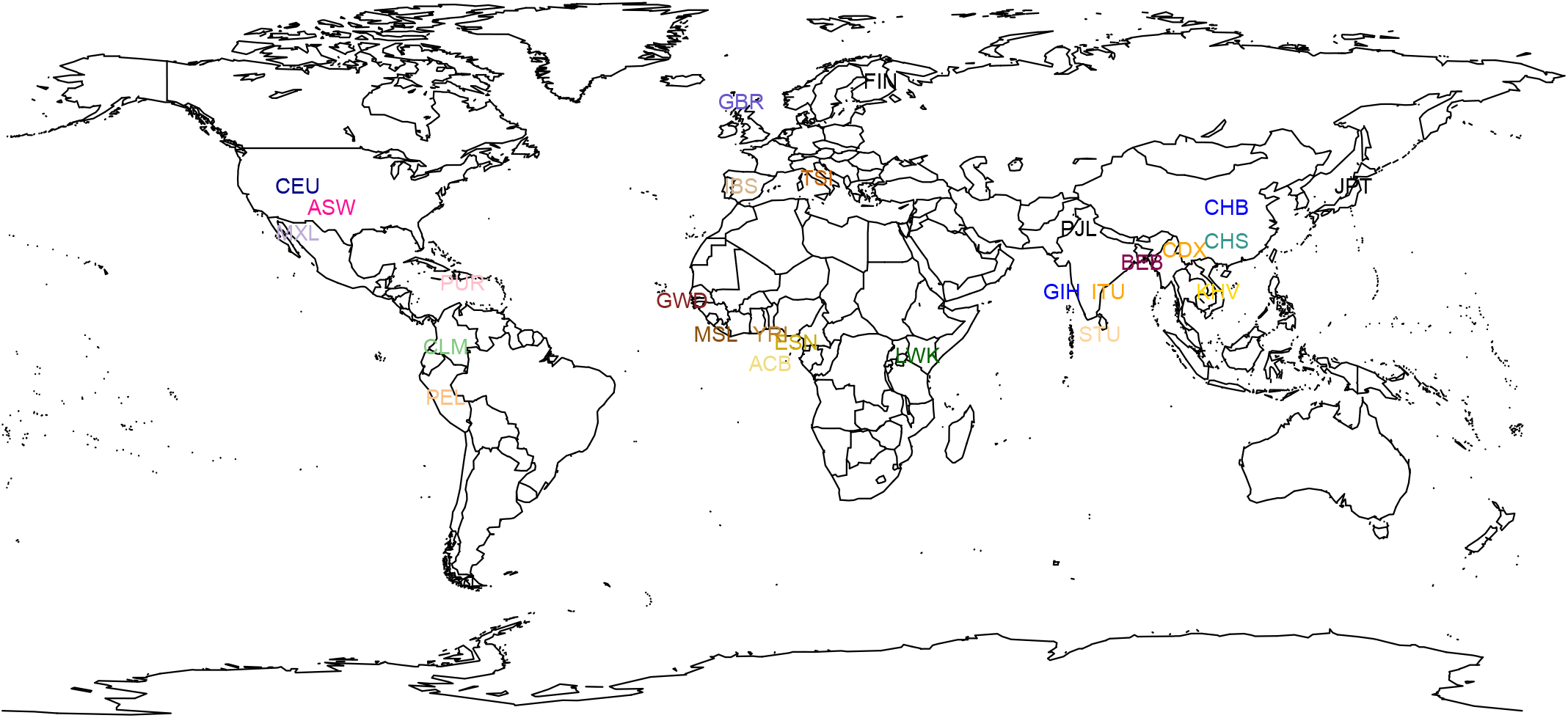
A world map where the location of the 26 populations are marked. Note these specific choices: the Utah Caucasian CEU is not moved to Europe, and southwest African American ASW is not moved to Africa; the African Caribbean in Barbados ACB is moved to the west coast of Africa; the Mexican-American in California MXL is moved to Mexico.

The lower-left cluster in Fig.6 contains 5 groups in East Asian: they are, in the rough ordering from north to south, Japanese in Tokyo (JPT), Han Chinese in Beijing (CHB), Han Chinese in South China (CHS), Chinese Dai in Xishuangbanna (CDX), and Kinh in Ho Chi Minh City, Vietnam (KHV). The change of color from light (south) to dark (north) can be clearly seen. However, the CHS points are more scattered

The upper-left cluster in Fig.6 is the South Asia (India subcontinent) populations, with this ordering in color lightness: Punjabi in Lahore,Pakistan (PJL), Gujarati Indian in Houston US (GIH), Bengali in Bangladesh (BEB), Indian Telugu in UK (ITU), and Sri Lankan Tamil in UK (STU). There is no question that PJL in the north, and STU/ITU in the south should be the two ends of the ordering. The GIH to BEB is a direction from west to east. Nevertheless, the trend shading change from top to bottom is clear.

The African populations (middle-left cluster) are mostly limited to the western Africa: Gambian in Western Gambia (GWD), Mende in Sierra Leone (MSL), Yoruba in Ibadan, Nigeria (YRI), Esan in Nigeria (ESN). The African Caribbean in Barbados (ACB) should also be of a western Africa origin. Only the Luhya in Webuye, Kenya (LWK) group is on the east coast. The origin of African-American in Southwest US (ASW) samples is not clear, so we use a distinctly different color. Interestingly, several ASW samples are closer to the America cluster, indicating a genetic admixture with native Americans. The west/east representatives of GWD and LWK do seem to anchor the two ends of the cluster in Fig.6.

The European populations are color ordered by this sequence: Finnish (FIN), Utah American (CEU), British (GBR), TSI for Toscani in Italy, and IBS for Iberian in Spain. Again, we observe a better separation between the two extremes: FIN at the north, and TSI/IBS at the south, with CEU and GBR more scattered.

America samples are the only one which do not form its own cluster, due to the well known admixture between indigenous America population and European colonists. Although we do not know the details of individual samples, most AMR samples do not seem to have a large Africa admixture (with a few exceptions). The Peruvian in Lima (PEL) samples are furthest away from the EUR cluster, which could be the center for a native America cluster. The Mexican-American in LA, US (MXL), Colombian in Medellin (CLM), Puerto Rican (PUR), appear to form a gradient extending from the PEL samples to the EUR samples.

Finally, a 3D t-SNE plot is shown in Fig.8. It illustrates both the continental separation in 4-5 large clusters, as well as sub-continental group patterns with gradient of colors present with each cluster as discussed above. In comparison, the PCA plot (not shown) has sub-populations belonging to the same continent tightly squeezed, making the separation among them difficult.

**Figure 8:**
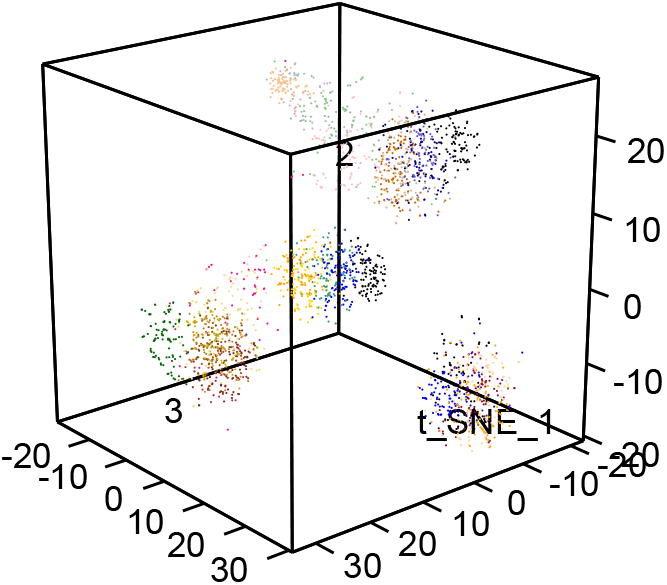
3-dimensional t-SNE which expands the plot in Fig.4. The cloud of dots on left is the AFR group, that on the right is SAS. The middle cluster is EAS, and the top cluster is ERU with a “tail” towards AMR.

## Discussion

Application of PCA to genetic data to display sample population stratification is a common practice [42], and the generic form of PCA plot is easily interpreted (e.g. Supplement Fig.S4 of [34]). However, using t-SNE is new. We show in Fig.1 that t-SNE can separate population from different continents as well as PCA.

Although some problems in the application of PCA to revealing population stratification remain in t-SNE, such as identification of the optimal number of SNPs to keep in the data in order to best show the population structure, whether rare variants should be used, etc, other problems are not an issue in t-SNE. For example, which high-order PCs should be kept to reveal the full population structure. In principle, one can use the proportion of variance explained to select the number of PCs. However, it is not clear what the threshold cutoff should be, and how to meaningfully interpret the contributions of a higher-order PC in visualization compared to its lower-order ones. We are less concerned about this issue with t-SNE, as t-SNE conducts dimension reduction by optimizing Kullback-Leibler distance between the raw data distributions, and a low-dimensional distribution in at most 3-dimension space, instead of via a PC selection.

The main feature of t-SNE, the ability to exhibit structures at multiple scales, is responsible for two observations in this paper. One is that an outlier in PCA does not appear to be an outlier in t-SNE. In other words, the display of the pattern in t-SNE is more robust against a small number of outliers. This property might be compared to robust PCA [43], nonlinear PCA [44, 45], local PCA [46] (note the meaning of “local” in this local PCA paper [39] refers to local genomic regions), or other robust methods in dimension reduction [47]. Another observation that both continental and sub-continental population structures can be viewed in a single t-SNE plot is also a consequence of this property of t-SNE. t-SNE is better than PCA at characterizing local structures, while equally well at preserving global structures.

The time computational complexity of standard PCA/SVD is *O*(*min*(*Np^2^* + *p*^3^, *pN*^2^ + *N*^3^) where *N* is the number of samples and *p* the number of factors [48]. If *N < p,* the computational complexity is *O*(*N*^3^). On the other hand, the computational complexity of t-SNE is *O*(*N*^2^) [28], which has an advantage over PCA. Of course, in specialized applications such as sparse matrix, approximate results, or partial results, the computational complexity can be improved [49, 50, 51].

In conclusion, though the current application of t-SNE in genomics is mostly limited to gene expression data such as those in single-cell RNA-seq, we found it quite useful in human genetic data, being able to display both global/continental scale and local/population scale patterns.

## Data and Methods

### 1000 Genomes Project data

The 1000 Genomes Project is a major effort to sequence the whole genome of more than 1000 samples [33, 34, 35]. The phase 3 1000 Genomes Project data for 2504 individuals was downloaded from *http://www.internationalgenome.org/data*.

### Genetic distance between samples

KING (Kinship-based INference for Genome-wide association studies), a computationally efficient program to calculate the person-person distance based on genetic data, was used: (*http://people.virginia.edu/∼wc9c/KING/*) [52].

### t-SNE

The R (*http://www.r-project.org/*) implementation of the t-SNE, *Rtsne* version 0.11 (June 30, 2016) *(https://cran.r-project.org/web/packages/Rtsne/*<u>or</u> *https://github.com/jkrijthe/Rtsne)* was used.

### PCA

The smartpca in the EIGENSOFT package [20] (v6.1.4) *https://data.broadinstitute.org/alkesgroup/EIGENSOFT/* was used for PCA of genotype data.

### Other programs

PLINK *http://pngu.mgh.harvard.edu/∼purcell/plink/* [53] was used for genotype file management and conversion. The 3D plot was generated by the *rgl* R package (*https://cran.r-project.org/web/packages/rgl/index.html*, 0.97.0). R (*http://www.r-project.org/*) statistical package was used for other general analysis and graphics.

## Acknowledgments

We would like to thank Andrew Shin for discussions. WL and JEC acknowledge the support from the Robert S. Boas Center for Genomics and Human Genetics, YY acknowledges the support from NSFC (11671375), and HH acknowledges the partial grant support from Fordam's IPGF.

